# Subjective value then confidence in human ventromedial prefrontal cortex

**DOI:** 10.1101/838276

**Authors:** Allison D. Shapiro, Scott T. Grafton

## Abstract

Two fundamental goals of decision making are to select actions that maximize rewards while minimizing costs and to have strong confidence in the accuracy of a judgment. Neural signatures of these two forms of value: the subjective value (SV) of choice alternatives and the value of the judgment (confidence), have both been observed in ventromedial prefrontal cortex (vmPFC). However, the relationship between these dual value signals and their relative time courses are unknown. We recorded fMRI while 28 men and women performed a two-phase Ap-Av task with mixed-outcomes of monetary rewards paired with painful shock stimuli. Neural responses were measured during offer valuation (offer phase) and choice valuation (commit phase) and analyzed with respect to observed decision outcomes, model-estimated SV and confidence. During the offer phase, vmPFC tracked SV and decision outcomes, but it not confidence. During the commit phase, vmPFC tracked confidence, computed as the quadratic extension of SV, but it bore no significant relationship with the offer valuation itself, nor the decision. In fact, vmPFC responses from the commit phase were selective for confidence even for rejected offers, wherein confidence and SV were inversely related. Conversely, activation of the cognitive control network, including within lateral prefrontal cortex (lPFC) and dorsal anterior cingulate cortex (dACC) was associated with ambivalence, during both the offer and commit phases. Taken together, our results reveal complementary representations in vmPFC during value-based decision making that temporally dissociate such that offer valuation (SV) emerges before decision valuation (confidence).

## Introduction

Every day, we navigate a maze of choices, guided by our subjective preferences and goals. When the route forks, multiple forms of value imbue the selection of one’s path. We assess the value of potential actions from their relative costs and benefits, as well as the value of our own judgment – accurate decisions are valuable decisions, regardless of the options.

Much progress has been made toward understanding how the brain resolves value-based decisions (1, 2). Ventromedial prefrontal cortex, in particular, is crucial for integrating reward and cost attributes of choice alternatives (3–6) and automatically tracking subjective value (SV)(7), which is the perceived utility of objective value information relative to the decision maker (7, 8). Neuroeconomic models benefit from mixed-outcome Approach-Avoidance (ApAv) tasks, which pose realistic, consequential choice scenarios (e.g. accept or reject offers of appetitive rewards contingent on aversive costs). Such fMRI studies in humans demonstrate multiple value representations in vmPFC including rewards, decision variables, SV, and the valuation models that inform SV (9–12).

Recent research suggests that vmPFC also signals choice confidence, how strongly one can believe they are making the best decision. For example, vmPFC activation varies with confidence about perceptual judgments (13–15) and lesions lead to atypical confidence reports on general knowledge tests (16) These findings could be evidence that vmPFC performs a valuation of one’s judgment, assigning high value to high confidence decisions (17). Confidence signals in vmPFC also accompany value-based decisions (17–19), which is intriguing given the relationship between confidence and value. Self-reported confidence takes a U-shaped function with respect to first-order valuation judgments (7, 20): decisions about extremely high- or low-value items elicit stronger confidence than decisions about items with neutral or ambiguous value. Similarly, vmPFC responses take a U-shaped function with respect to value in risky decision making (21). Accordingly, Lebreton et al. (17) operationalized confidence as the quadratic extension of value and elegantly demonstrated that vmPFC tracks modeled confidence, even in the absence of explicit ratings.

Given this literature, vmPFC should track both SV and its quadratic extension, confidence, in mixed-outcome ApAv decision making, but this has not been explicitly tested. Specifically, confidence about accept choices (typically positive SV) should increase as SV increases whereas confidence about reject choices (typically negative SV) should increase as SV decreases. It is unknown how vmPFC represents both SV and confidence in value-based choices, particularly when confidence and value are inversely related (i.e. reject choices). One possibility is that confidence evolves in parallel with decision variables (22). Early confidence-related signals have been recorded from frontal and parietal sites (23, 24), including vmPFC (7,14,19). Alternatively, confidence may evolve later through retrospective metacognitive judgments or continued deliberation after choice commitment (25–29), both of which can recruit medial prefrontal cortex (30–32).

To test this, we deployed a two-phase ApAv task in which participants accepted or rejected offers of monetary rewards paired with painful shock stimuli. BOLD responses were measured during offer valuation (offer phase) and choice valuation (commit phase). Observed decision outcomes and model-based estimates of SV and confidence were used to predict neural activity in vmPFC and elsewhere during both phases of decision making.

## Methods

### Experimental Design

#### Participants

We report the data from 28 paid volunteers that participated in the study (17 women, 27 right-handed, mean age = 21.9, sd=2.8). One additional participant completed the study but was removed from analyses due to significant susceptibility artifacts causing excessive errors in spatial normalization. No participants had a history of neurological injuries or illnesses or current daily use of psychoactive medications. All participants provided written consent in accordance with the Institutional Review Board at the University of California, Santa Barbara.

#### Session overview

All testing was performed on the same day. Participants first provided informed consent and were screened for disqualifying criteria. Because our task entailed ApAv decisions with pain stimuli, participants next underwent a pain thresholding procedure and were familiarized with the mapping between experimental stimuli and the pain intensities they represented. Then, participants performed the decision making task while in the MRI scanner. Finally, after they had completed all experimental tasks and been removed from the scanner, participants received the pain stimuli and monetary rewards associated with the choices they made during the task.

#### Pain thresholding procedure

We used mild cutaneous electrical shocks as pain stimuli. Pain thresholding allowed us to control for individual differences in pain tolerance for electrical shocks such that a given experimental stimulus was associated with the same subjective experience of pain across participants. To identify participant-specific minimum and maximum shock intensities, participants completed a pain thresholding procedure before the decision-making task, outside of the MRI scanner. Electrical shocks were administered with a constant current stimulator (Digitimer DS7A, Digitimer, Great Britain) controlled by a train generator (DG2A Train/Delay Generator, Digitimer, Great Britain). Each shock had a duration of 1 s and a frequency of 100 Hz with a 2 ms waveform. Two adhesive electrodes were placed on the back of the participant’s hand approximately 1 inch above the wrist and connected to the stimulator. When a shock was administered, electric current was run between the two electrodes, causing an aversive sensation that is increasingly painful at higher levels of current.

The first stage of thresholding was a ramp-up procedure in which the experimenter delivered several shocks, each time increasing the intensity by 1 mV. The participant was instructed to report three thresholds: the lowest intensity at which they detected the shock, the lowest intensity at which the shock caused discomfort, and the intensity at which the shock became unbearably painful. The second stage was a rating procedure in which the participant received fourteen shocks with intensities between the discomfort threshold and unbearable threshold and rated the pain from each on a 0-10 scale. Next, both the ramping and rating stages were repeated to verify that we had accurately identified the participant’s pain tolerance. Finally, the pain ratings and shock intensities from the second rating procedure were fit with a sigmoid function to model the relationship between shock intensity and perceived pain for this individual (33). Based on this sigmoid function, we identified the shock intensity that predicted a pain rating of 8 out of 10. No payout trials during the decision task had costs exceeding this value.

#### Familiarization Procedure

Before the decision making task, participants were shown five example offer stimuli illustrating costs of 5%, 25%, 50%, 75% and 95% while the experimenter delivered a shock at the corresponding intensity relative to their discomfort and maximum pain thresholds (with 0% = minimal discomfort and 100% = maximal pain). Participants were instructed to remember the sensation associated with the example shocks and use these as points of reference when making choices during the task, but that the real offers would include shocks ranging anywhere between 0 and 100% intensity, not just at the example levels.

### ApAv Task

#### Offer stimuli

We adapted an approach–avoidance task similar that implemented in non-human primates by Amemori and Graybiel (4, 34) in which participants accepted or rejected offers with mixed appetitive and aversive outcomes. On each trial, the participant was offered a certain amount of money in exchange for receiving a shock of a certain intensity (Fig 1a) . The participant chose either to accept *both* the money and the shock or reject both (i.e. receiving a monetary reward was always contingent on also receiving its associated cost). Therefore, trial stimuli were deterministic mixed-outcome choice scenarios as there were no manipulations involving probabilistic chances or risks of receiving the shock and/or money that could be used to form decision strategies.

**Fig 1:**
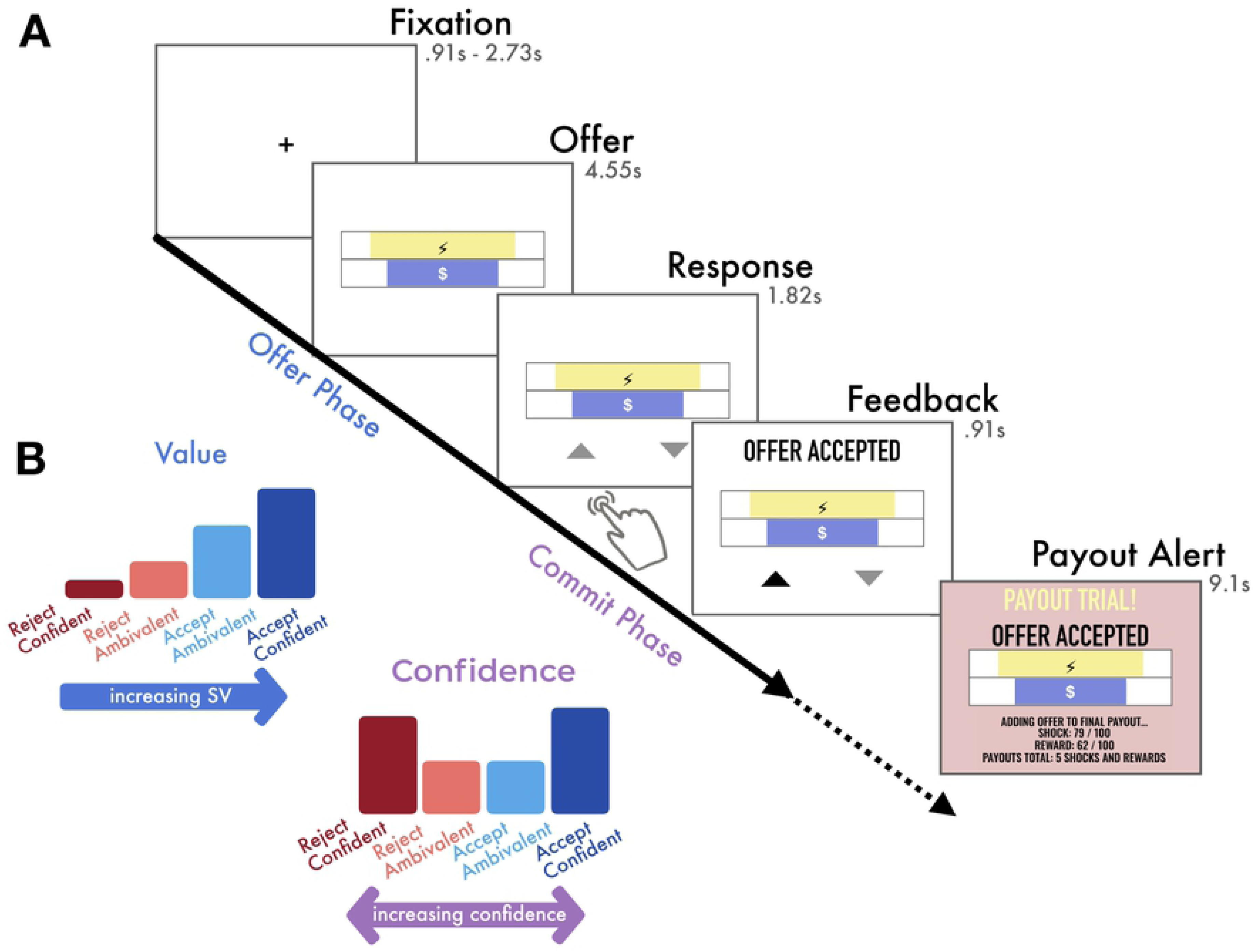
Experimental task and variables of interest. **(A)** Experimental Task: On each trial, the participant was offered one monetary reward ($ bar width represents amount, ranging continuously $0.01-$1.50) contingent on enduring 1 painful shock (lightning bolt bar width represents pain intensity, ranging continuously from minimally to maximally painful). Participants were instructed to use the offer phase to evaluate the offer and to decide if they would accept or reject it, but they were not yet able to respond. During the commit phase, response mappings appeared and participants made a left or right button press according to the location of the triangle representing their choice (up triangle = accept, down triangle = reject), which varied randomly between trials to prevent preparation of motor responses during the offer phase. After submitting a response, the corresponding triangle was highlighted. Finally, feedback indicating whether the offer was accepted or rejected was added. On payout trials (10 random trials of 189 total), a payout alert followed the feedback. If the participant had accepted the offer, they would receive the monetary reward and also endure the shock at the end of the task, otherwise the participant would receive neither. **(B)** Illustration of variables of interest: We were interested in two types of value inherent in economic decision making: the perceived value of the offer stimulus (SV) accounting for its cost and reward attributes, and the value of one’s judgment (confidence), which measures the extent to which one believes they are making the best decision. For accept decisions (blue bars), there is a positive relationship between SV and confidence: as SV increases, one becomes increasingly confident that accepting the offer is the best decision. However, for reject decisions (red bars), there is an inverse relationship between SV and confidence: one becomes increasingly confident about rejecting offers as SV *decreases.* We tested whether they are evaluated simultaneously or during different phases of decision making.

The offer stimuli were two horizontal bars, one had an overlaid $ symbol and illustrated the offered reward and the other had an overlaid lightning bolt symbol and illustrated the contingent cost. The width of the bars represented the amount of each attribute being offered on the current trial, with monetary rewards ranging continuously from $0.01 to $1.50 and shock intensities ranging continuously from minimal discomfort to maximal pain. Each bar was within a larger rectangular frame that illustrated the maximum possible bar width. The bars were blue and yellow, and color-attribute mappings varied between participants. The relative position of the cost and reward bars (i.e. which bar was above the other) alternated between blocks. The offer stimuli were centrally aligned and stacked just above and just below the vertical midpoint of the display. Overall, there were 189 offers, each presented as a single trial, unless the participant failed to respond, in which case the offer was repeated at the end of the experiment. Trials were split between six functional runs, with each run containing 31 or 32 trials. All participants viewed the same set of offers, presented in the same pseudo-random order that systematically covered all quadrants of the decision space.

#### Decision making task

On each trial, the participant first saw a fixation point in the center of the screen for either 910 ms, 1820 ms or 2730 ms, randomly varied between trials. Then, the offer stimulus was shown for 4550 ms. The participant was instructed to use this time to evaluate the offer but could not yet indicate a choice. Next, the offer remained on the screen and two response mappings appeared in left and right positions beneath the offer stimulus for 1820 ms. During this time, the participant was required to respond with a button press, indicating their commitment to either accept (approach) or reject (avoid) the offer. All stimuli remained on the screen for the remainder of the response interval, and after the response was submitted, the response mapping corresponding to their choice was highlighted. Participants were allowed to change their response within the 1820 ms response interval. The response mappings were an upward pointing triangle representing the accept option and a downward pointing triangle representing the reject option. The participant used either the index or middle finger of their right hand to press either the left or right (respectively) button on a Cedrus LP-RH response pad transmitting through a Lumina LSC-400 controller (Lumina, Cedrus Corportation, San Pedro, CA, USA), according to the location of their preferred choice option. The left/right positions of the choice options varied from trial to trial, preventing the participant from pre-planning a motor response before the response phase. Finally, a 910 ms feedback interval followed the response interval. Feedback included the offer stimulus, which remained on the display, and text stating whether the offer was accepted or rejected. On payout trials, a small subset of all trials, an additional 9100 ms payout alert followed the decision feedback.

Because we were interested in value-processing during different phases of decision making, our analyses separately analyzed neural responses from the offer phase and the commit phase of each trial. The offer phase was the 4550 ms in which the offer was displayed but the participant was not yet able to submit a response. The commit phase was the 2730 ms comprising the response interval (1820 ms) and the feedback interval (910 ms).

#### Payouts

Participants did not receive the rewards and shocks for all accepted trials. Instead, during trial generation, 10 pseudorandomly selected offers were tagged as payout trials. On these trials, if the participant accepted the offer the payout alert indicated that they would receive both the monetary reward and the shock. If they rejected the offer, the payout alert indicated they would receive neither. The payout alert also showed the cumulative number of payouts accepted so far. Participants were encouraged to assume that every trial was a payout offer and decide to accept or reject it accordingly, as they would not see the payout alert until after they’d submitted their choice. Although participants were notified whether an offer was a payout immediately after choosing to accept or reject it, the actual delivery of monetary rewards and shocks from payout trials occurred after they had completed the entire decision making task and had been removed from the MRI scanner. In pilot studies, we found that providing realtime payout feedback, relative to showing which trials were payouts at the end of a fixed-length block or at the end of the task, reduced the proportion of offers that were accepted. We believe that this may indicate that instant feedback reduces temporal discounting of future costs, relative to when that information is delayed, even though in both cases the time that the costs were actually delivered was the same (after completing the entire task).

All participants also performed an additional cost-benefit decision making task after they completed the approach-avoidance task, not reported here. Twenty-four of the participants also underwent simultaneous physiological recordings of impedance cardiography while performing the decision making tasks in the MRI scanner, impedance cardiography data is not included here and will be presented elsewhere.

### Behavioral statistical analysis

#### Estimating SV from decision behavior

The subjective value (SV) of each offer can be represented as

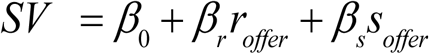

where *r_offer_* is the available reward, *S_offer_* is the contingent shock, *β_r_* and *β_s_* describe how strongly the individual subject weights rewards and shocks, and *β*_0_ is the individual’s intercept, indicating their intrinsic motivations to pursue reward versus avoid shock (33). We separately modeled each participant’s choice data with logistic regression, a specialized from of the generalized linear model, using the glm package in R. Offers to which participants failed to respond before the decision deadline were repeated at the end of the task. Choice outcomes from the second presentation were included in the dataset used for generating valuation models but these trials were excluded from the fMRI analysis. Participants’ model estimates were used inversely to predict the perceived subjective value of each offer, SV, and the likelihood that it would be accepted, P(Acc) specific to each individual on each trial.

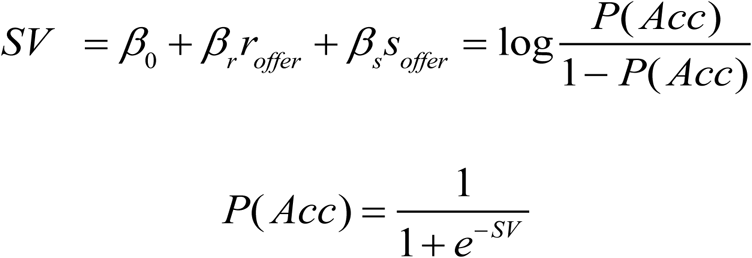

#### Estimating confidence from SV

Here, we operationalize choice confidence as the extent to which one can believe they are making the best decision. In our task, given the options to either accept or reject, the best choice depends on the perceived value of the offer and consequently choice confidence is maximized when SV unambiguously points to one choice over the other. Therefore, confidence varies in a U-shaped function with respect to increasing SV such that when SV is extremely high or extremely low, one can be highly confident in choices to accept or reject the offer, respectively, but when an offer’s SV is neutral there is more ambivalence about the decision (Fig 1b). We quantify confidence in accordance with Lebreton et al. (2015), who demonstrated that decision confidence can be estimated from the quadratic extension of perceived value in the absence of explicit confidence ratings.

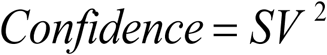

We used these trial-by-trial estimates to test whether behavioral response times and neural activity varied with respect to choice confidence, see below. Notably, because we did not collect metacognitive self-reported confidence ratings, we do not intend to make explicit claims about the subjective experience of confidence. Previous research has found mixed results regarding how closely model-estimated confidence varies with self-reported confidence, see the discussion for a further explanation. Therefore, we primarily aim to describe neural responses that vary with model-estimated confidence.

We also inspected the relationship between confidence and choice *outcomes* by categorizing trials into confident accepts (AccCon), ambivalent accepts (AccAmb), confident rejects (RejCon), and ambivalent rejects (RejAmb). Each participant’s trials were binned by choice outcome and then we performed a median split on *SV* for both choice bins. Accepted offers in the upper 50% SV were assigned to AccCon and the rest were assigned to AccAmb. Rejected offers in the upper 50% SV were assigned to RejAmb, and the rest were assigned to RejCer. Notably, because we observed substantial individual differences in subjective valuation, both parameterizing and categorizing the offers according to individual-specific model-based estimates allowed us to describe objectively identical offers differently for each participant, according to their own individual preferences. This was a critical feature of our study that allowed us to observe group-wide neural responses that were specific to decision outcome, SV, and confidence, regardless of the objective properties of the stimulus.

In our paradigm, P(Acc) varies as a sigmoidal function of SV while choice confidence varies as a U-shaped function of SV. Consequently, when P(Acc) is close to either 0 or 1 there is high choice confidence in either accepting or rejecting the offer, respectively and P(Acc) is close to .5, there is high ambivalence about committing to either choice. In a two-dimensional decision space in which reward values are represented on one dimension, pain values are represented on the other, and the modeled decision boundary is the vector along which P(Acc) = .5, the choice confidence associated with a reward-pain offer pair increases with its distance from the decision boundary. We present individual choice outcomes and binned choice confidence overlaid on estimated P(Acc) throughout the decision space.

### Neuroimaging Data Acquisition and Preprocessing

#### Neuroanatomical ROI

We investigated task-related involvement within an a priori region of interest (ROI), vmPFC. The ROI was anatomically defined from the Harvard-Oxford cortical structural probabilistic atlas (https://fsl.fmrib.ox.ac.uk/fsl/fslwiki/Atlases), and includes all voxels with at least 25% likelihood of being located within areas labelled Frontal Medial Cortex or Subcallosal Cortex.

#### MRI protocols

Anatomical and functional MRI data were collected on a Siemens 3T Magnetom Prisma Fit with a 64-channel phased-array head and neck coil (58 channels active for functional coronal imaging). High-resolution 0.94 mm isotropic T1-(TR=2500 ms, TE=2.2 ms, FA=7°, FOV=241 mm) and T2*-weighted (TR=3200 ms, TE=570 ms, FOV=241 mm) sagittal sequence images were acquired of the whole brain. Next, functional MRI recordings were collected while participants performed the decision making task. For each functional run, a multiband T2*-weighted echo planar gradient-echo imaging sequence sensitive to BOLD contrast was acquired (TR=910 ms, TE=32 ms, FA=52°, FOV=192 mm, multiband factor 4) provided by the Center for Magnetic Resonance Research in accordance with a current license. Each functional image consisted of 64 coronal slices acquired perpendicular to the AC-PC plane (3 mm thick; 3×3 mm in-plane resolution). Coronal orientation is necessary when acquiring simultaneous impedance cardiography to avoid artifact (35).

#### MRI pre-processing

Anatomical data was skull-stripped using Advanced Neuroimaging Tools (ANTs) brain extraction script (36). All other image pre-processing was performed with FMRIB’s Software Library (FSL, www.fmrib.ox.ac.uk/fsl). The first 10 volumes of each functional run were removed to eliminate non-equilibrium effects of magnetization occurring before the start of the task. The remaining functional volumes were skull-stripped using BET (37) motion corrected using MCFLIRT (38), spatially smoothed using a Gaussian kernel of FWHM 5mm, intensity normalized relative to the grand-mean of the entire 4D dataset by a single multiplicative factor, and underwent high-pass temporal filtering (Gaussian-weighted least-squares straight line fitting, with sigma=50.0s).

In preparation for group analyses, participants’ six functional runs were registered to their anatomical image and then to the Montreal Neurological Institute (MNI) 2mm averaged 152-brain template included with FSL distributions, using FSL’s linear image registration tool with 12 degrees of freedom (FLIRT; (38, 39). Refinement of the latter transformation was carried out with FSL’s nonlinear registration image registration tool (FNIRT) with a 10mm warp resolution (40, 41).

### Neuroimaging Statistical Analysis

#### fMRI analysis

FMRI data processing was carried out using FEAT (FMRI Expert Analysis Tool) Version 6.00, part of FSL (FMRIB’s Software Library, www.fmrib.ox.ac.uk/fsl). Time-series statistical analysis was carried out using FSL’s improved linear model (FILM) with local autocorrelation correction (42). We performed whole-brain statistical analyses with two general linear models (GLMs), as described below. Both GLMs had separate terms for the offer phase (off) and the choice commitment phase (com) of each trial. Offer phase regressors were time-locked to the onset of the offer and had a duration of 4.55s, during which the participant assesses the SV of the offer but cannot yet respond. Commit phase regressors were time-locked to the onset of the response mappings and had a duration of 2.73 seconds, during which the participant submits their decision about the offer and then views feedback confirming their choice. Finally, the GLMs also included a nuisance regressor for payout notifications, which were not used in analyses of interest but were intended to absorb variance in neural responses associated with subjective value (as payout notifications included an image of the payout offer) but unrelated to decision making processes. The payout regressor was time-locked to the onset of the payout notification and had a duration of 4.55 s, and the remaining 4.55 s of the payout notification screen was included with baseline activity.

Both analyses were performed at three sequential levels. First, at the run level, each participant’s six runs were separately modeled to find mean within-run activity corresponding each regressor and contrast images were generated by estimating pairwise differences between conditions. Then, at the participant level, run-level data was combined (fixed effects) to find the participant’s overall mean response relating to each regressor and contrast. Finally, at the group level, the participant data was combined (mixed-effects treating participant as a random effect with FSL’s FLAME 1) to find the group-wide mean responses for each regressor and contrast.

We tested the results of each contrast with non-parametric permutation testing at the whole brain level with threshold-free cluster enhancement (TFCE), implemented with FSL’s Randomise. This approach minimizes false positives by deriving a null distribution from the voxelwise data rather than assuming a parametric null distribution. For one-sample t tests, the distribution is created by iteratively multiplying statistical map values by 1 or -1, we performed 5,000 permutations of each contrast. TFCE detects clusters of contiguous voxels without setting an arbitrary cutoff for minimum cluster size or voxel statistic but rather summarizes the cluster-wise evidence at each voxel, against several types of cluster-forming thresholds and controls the family-wise error (FWE) rate at p=.05 (43, 44). We present figures with voxel-wise T-values from all voxels that survived whole-brain TFCE correction. Some contrasts yielded significant voxels across contiguous but widespread regions of cortex, and consequently reporting only the peak voxel of a cluster would obscure other local maxima in different anatomical regions. Therefore, we also report the coordinates, t statistics, Brodmann area, and anatomical structure labels from Automated Anatomical Labelling the of the MNI atlas for local maxima within each cluster in Table S1. Local maxima were found with the cluster command provided with FSL and labelled with label4MRI, a freely available toolbox for R (https://github.com/yunshiuan/label4MRI). We additionally report results from sub-conditions of GLM2 restricted to and TFCE corrected only within our primary ROI of interest, vmPFC.

**Table 1:** Comparison of linear mixed models predicting vmPFC responses from value. Results of model comparison are depicted in Table 1. The first column identifies the model and variables specified as fixed effects from that model. Columns 2-6 give the model estimate, 95% confidence interval (95CI), and t statistic and p value estimated from Satterthwaite approximation. Column 7 is the standard deviation (SD) of the random effect (Intercept | Subject). Columns 8-11 are overall model statistics: variance explained by fixed effects (Marg. = marginal R^2^), variance explained by fixed and random effects together (Cond. = conditional R^2^), and overall model AIC, and BIC. The final two columns are with χ2 statistics and p values from likelihood ratio tests (after refitting models with maximum likelihood estimates), describing improvement of fit from adding quadratic term to the original linear model.

|  | Fixed Effects |  |  |  |  | Random Effect<br>*SD* | Model Fit |  |  |  | Model Comparison |  |
| --- | --- | --- | --- | --- | --- | --- | --- | --- | --- | --- | --- | --- |
| | Estimate | 95CI | *t* | *p* | | | Marg. $R^2$ | Cond. $R^2$ | AIC | BIC | $\chi^2$ | *p* |
| Offer: Linear Model |  |  |  |  |  | 14.58 | 0.05 | 0.70 | 901.1 | 912.0 |  |  |
| Intercept | -19.08 | -24.74 -13.28 | -6.56 | <0.001 |  |  |  |  |  |  |  |  |
| P(accept)-P(reject) | 5.10 | 2.73 7.41 | 4.38 | <0.001 |  |  |  |  |  |  |  |  |
| Offer: Quadratic model |  |  |  |  |  | 14.57 | 0.05 | 0.70 | 903.1 | 916.7 | 0.002 | 0.966 |
| Intercept | -19.00 | -25.78 -12.20 | -5.50 | <0.001 |  |  |  |  |  |  |  |  |
| P(accept)-P(reject) | 5.11 | 2.76 7.40 | 4.33 | <0.001 |  |  |  |  |  |  |  |  |
| (P(accept)-P(reject))<sup>2</sup> | -0.12 | -5.76 5.44 | -0.04 | 0.967 |  |  |  |  |  |  |  |  |
| Commit: Linear Model |  |  |  |  |  | 19.17 | 0.01 | 0.55 | 1014.5 | 1025.3 |  |  |
| Intercept | -9.07 | -17.08 -1.02 | -2.28 | 0.031 |  |  |  |  |  |  |  |  |
| P(accept)-P(reject) | 3.63 | -0.48 7.78 | 1.76 | 0.082 |  |  |  |  |  |  |  |  |
| Commit: Quadratic model |  |  |  |  |  | 19.18 | 0.07 | 0.61 | 1002.0 | 1015.6 | 14.43 | < 0.001 |
| Intercept | -20.86 | -30.57 -10.96 | -4.20 | <0.001 |  |  |  |  |  |  |  |  |
| P(accept)-P(reject) | 2.87 | -0.96 6.63 | 1.49 | 0.140 |  |  |  |  |  |  |  |  |
| (P(accept)-P(reject))<sup>2</sup> | 18.16 | 9.21 27.27 | 3.91 | <0.001 |  |  |  |  |  |  |  |  |

### GLM1: Parametric analysis SV and confidence

GLM1 was a parametric statistical analysis to observe neural activity modulated by SV and confidence (CD) during the offer and commit phases of each trial:

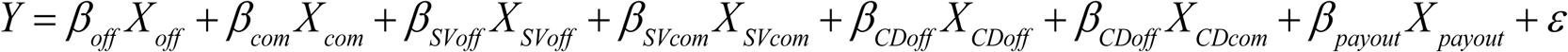

where Y is the time series of a given voxel predicted by a design matrix with one row for each time sample and one column for each of 7 trial regressors, convolved with a canonical gamma hemodynamic response function. The model included categorical terms (*β_off_X_off_, β_com_X_off_*) to isolate task related activity during the offer and commit trial phases (onsets and offsets described above). Parametric terms *β_SVoff_X_SVoff_* and *β_CDoff_X_CDoff_* were orthogonalized with respect to *β_off_X_off_* to capture variance in neural activity during the offer phase explained by trial-by-trial SV and confidence (for which regressors were range normalized by z-score). Likewise, *β_SVcom_X_SVcom_* and *β_CDcom_X_CDcom_* were orthogonalized with respect to *β_com_X_com_* and modeled neural activity related to SV and confidence during the commit phase.

For the purposes of visualizing gradual changes in neural responses over the course of the trial, GLM1 was re-estimated with a finite impulse response (FIR) model. Rather than splitting the trial into two phases, neural responses tracking SV and confidence were measured at each TR (910 ms per TR) beginning at the onset of the offer stimulus. The model was identical to GLM1 except for the following modifications. Instead of two categorical regressors for the offer and payout outsets there was a single categorical regressor for the offer stimulus onset and instead of two parametric terms for both SV and confidence there was only one for each. Instead of convolving regressors with a gamma HRF, FIR basis functions sampled neural responses at each of 18 discrete time points, with the first sample taken 0 s after trial onset and the last sample taken 16.38 s after offer onset. Given the delay of the BOLD response, the 16 second window was intended to capture the full progression of neural responses that began at some point between offer onset and the end of the trial. FIR basis functions were fit for each of the trial regressors (categorical offer regressor, parametric SV regressor, and parametric confidence regressor) as well as the payout nuisance regressor, for which the appearance of the payout notification marked the stimulus onset. As in GLM1, these analyses were first performed at the run level and then combined at the participant level, both of which were conducted in FSL’s FEAT. Participants’ mean parameter estimates for SV and confidence from each of the 18 FIR times were extracted from the anatomical vmPFC ROI and used for visualization.

### GLM2: Categorical choice by confidence analysis

GLM2 estimated categorical variance in neural responses during the offer and commit phases of each trial:

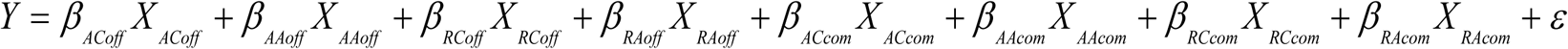

The first four terms modeled the offer phase of confident accepts, ambivalent accepts, confident rejects, and confident accepts, respectively. The next four terms modeled the commit phase of the same conditions. The ninth regressor modeled payout notifications. Our contrasts compared each condition with baseline, tested main effects of choice outcome (i.e. accepted offers vs. rejected offers irrespective of choice confidence) and choice confidence (i.e. high confidence choices vs. low confidence choices irrespective of choice outcome) separately during the offer and commit phases, as well as select pairwise comparisons within these conditions (AccCon vs. AccAmb, and RejCon vs. RejAmb), which were specifically inspected within our anatomical ROI, vmPFC.

#### Phase-specific responses in vmPFC

To further characterize the pattern of vmPFC responses during the offer and commit phases, we extracted mean parameter estimates within vmPFC for each condition at each trial phase. We tested whether vmPFC simultaneously tracks value and confidence, or if these two signals modulate vmPFC responses during different phases of decision making. Following the logic of Lebreton et al., (2015), we assumed that the function that decision phase-specific vmPFC responses take across trial conditions of ascending value (RejCon < RejAmb < AccAmb < AccCon) would be indicative of the information being processed during that trial phase. A linear increase of vmPFC response magnitudes would be associated with value processing, whereas a quadratic function would be associated with confidence processing. If vmPFC simultaneously tracked confidence and value, model terms for both the linear and quadratic extensions of value would be necessary to fully explain the observed pattern of vmPFC activity.

To tailor the predictors in these models to individual participants, we used the mean perceived value across trials from each condition, calculated separately for each participant. SV varies substantially between participants depending on the consistency of their choice behavior. Specifically, slight differences in participants’ choice consistency lead to substantial differences between their model-estimated SV predictors. Consequently, there was a large range of SV and SV^2^ throughout the group and many participants’ data only spanned only a portion of that range, making it difficult to draw conclusions at the group level. P(Acc) and P(Rej) (the latter is equivalent to 1-P(Acc)) are always restricted to the range of 0 to 1. Consequently, P(Acc)-P(Rej) normalizes value to range from -1 to 1 while preserving the sign of SV, with negative values predicting reject choices, positive values predicting accept choices, and values surrounding zero indicating decision ambivalence. Conversely, normalizing SV at the individual level (e.g. z-score) would not preserve the sign of raw SV. Furthermore, whereas in GLM1 we observed parametric modulation by SV, in GLM2 trials were binned according to observed choice, which is more specifically related to P(Acc)-P(Rej). Therefore, we used P(Acc)-P(Rej) as value predictors in our ROI analysis, which improved consistency for groupwide analysis while retaining the sign of SV. Previous research has used a similar approach (17).

Each participant’s mean vmPFC parameter estimates from the four trial conditions were predicted as a function of their mean P(Acc)-P(Rej) of that condition using linear mixed effects regression, implemented with the lme4 package for R (45).

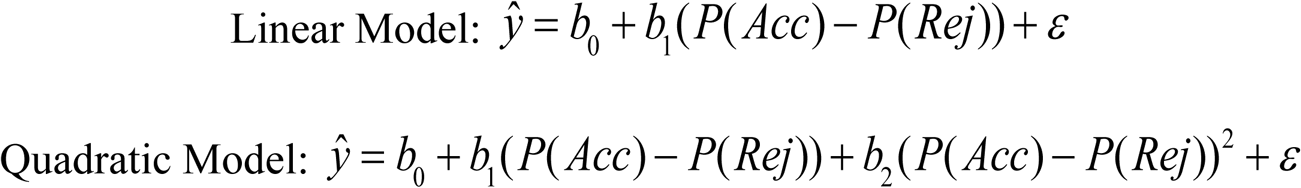

The linear and quadratic extensions of P(Acc)-P(Rej) were specified as fixed effects and we included a random effect on the model intercept across subjects to account for baseline variation in vmPFC parameter estimates. Predictor values entered into the model were participants’ mean P(Acc)-P(Rej) of trials from each of the four conditions. Both models were separately fit to parameter estimates of BOLD responses in vmPFC during the offer phase and during the commit phase. We report model fits as well as the results of model comparisons. Due to ambiguity in estimating denominator degrees of freedom, linear mixed model fits are not best evaluated by p-values. However, significance can be inferred from confidence intervals constructed by iteratively sampling the model posterior to estimate the likelihood of the observed parameter estimates. We ran 5,000 simulations using the posterior distributions over each parameter from the mixed models using the merTools package for R. We report significant parameters with 95% CIs that do not span zero. For interested readers, corresponding p-values estimated with Satterthwaite’s method implemented in the lmerTest package for R are also provided. Model comparison (ordinary likelihood ratio test) and relative AIC and BIC values were used to determine the best fitting model for the offer phase and for the commit phase.

### Behavioral correlates of SV and confidence

We estimated confidence as the quadratic extension of value in accordance with a similar study that validated this operationalization by demonstrating that both response times (RTs) and self-reported confidence were related by the inverse quadratic to value, and thus RTs and confidence were negatively correlated. Notably, that study found that quadratic extension of value (i.e. model-based confidence) better predicted self-reported confidence than RTs, suggesting that while RTs were a useful behavioral correlate of subjective confidence, they didn’t fully explain variance in metacognitive confidence ratings (7, 17).

We aimed to measure neural correlates of implicit, naturalistic experiences of confidence during decision making. Therefore, our task did not solicit metacognitive confidence ratings. Instead, we used model-based confidence (estimated in accordance to previous literature) to predict changes in BOLD responses. Given previous findings that model-based confidence better predicted confidence ratings than RTs, we did not take RTs to be a direct proxy for the subjective experience of confidence. Nonetheless, it was important to verify that model-estimated confidence mad a meaningful relationship with behavior in our task, that is, to the time it took participants to commit to a decision. Specifically, an inverse quadratic relationship between behavioral RTs and model-estimated confidence would indicate that it took participants longer to commit to decisions that were associated with lower degrees of model-estimated confidence. To test this, we fit RTs with mixed effects regression (lme4 package for R; (45)). Fixed effects specified the linear and quadratic extensions of value and a random effect on the model intercept was included to account for baseline variation in RT.

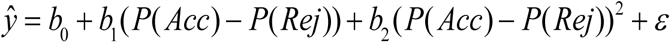

Value predictors entered into the model were generated by sorting model-estimated value into 5 equally spaced bins and the dependent measures were participants’ mean RTs for each bin. The model was tested again using z-scored SV as value predictors to verify that relationship between value and RTs were consistent regardless of the method used to estimate value.

## Results

### Behavioral choice models and RTs

A separate logistic regression model was fit to each participant’s decisions to accept or reject shock/reward offers made while undergoing fMRI. All participants’ model fits had significantly positive reward coefficients and significantly negative shock coefficients, indicating that both offer attributes influenced SV in the intended direction, despite individual differences in their relative contribution to choice outcomes (β_r_ estimates: mean= .199, range = [0.064, 0.547], p values all <.001), costs (β_s_ estimates: mean = -0.170, range = [-0.547, -0.050], p values all <.001) (Fig 2). Moreover, there was great variation in participants’ model intercepts with ranges that spanned zero, suggesting strong individual differences in baseline tendencies to accept or reject offers (β_0_ estimates: mean = 1.057, range = [-6.420, 9.750]). On average, participants tended to accept more offers than they rejected (mean=61.3%, sd=17.8%), and seemed to be engaged in the task (98.2% of all trials received responses before the 1.8s decision deadline). Finally, linear mixed effects regression of behavioral RTs confirmed that RTs were quadratically related to SV and negatively correlated with estimated confidence, signifying that choices with low estimated confidence indeed took longer. This relationship between RT and confidence is consistent with previous research that defined confidence in a similar form (17).

**Fig 2:**
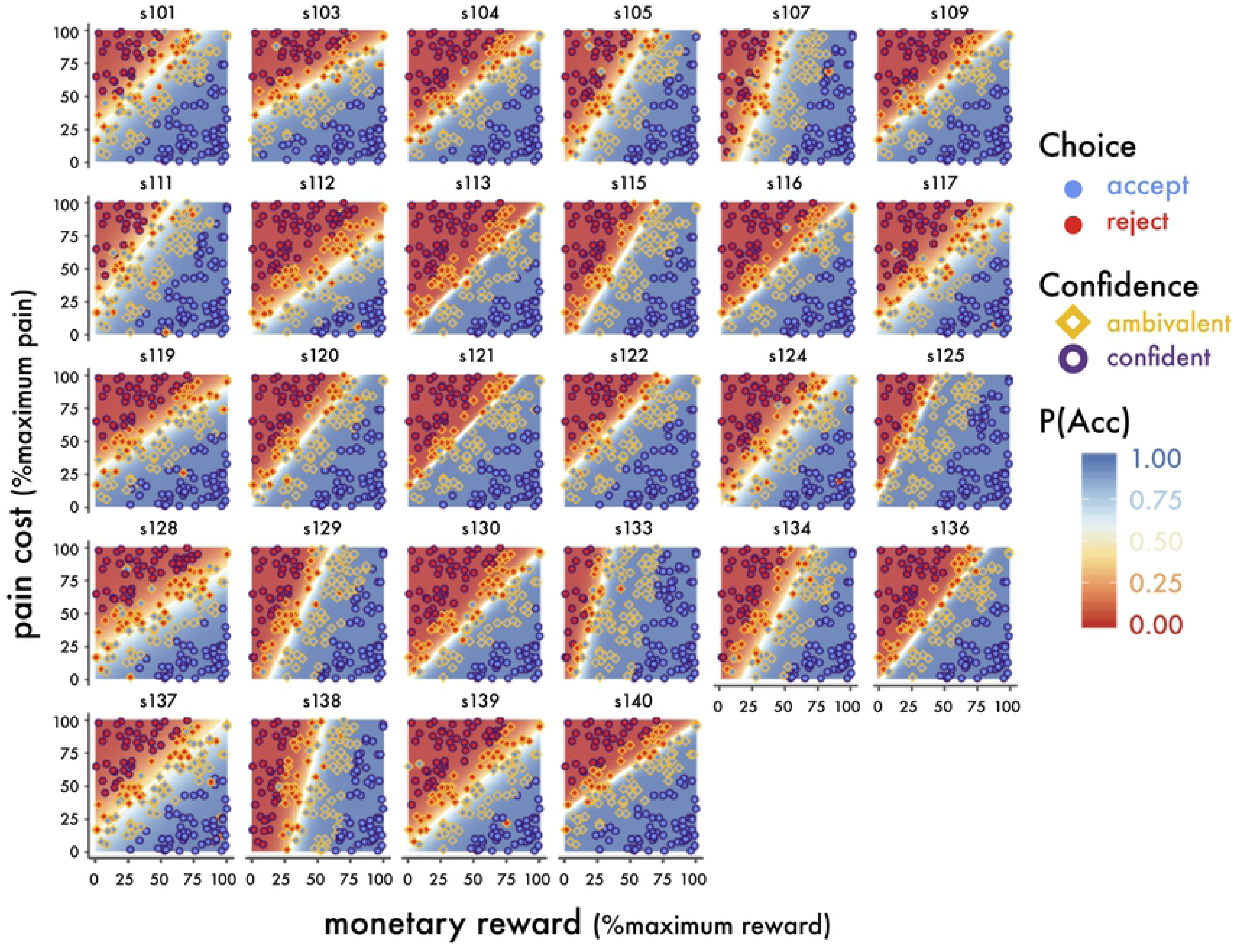
Observed decisions and model-based estimates of confidence and value. Individuals’ choice outcomes are plotted over the decision space, which is shaded according to model-based P(Acc). Each coordinate is a possible offer, with the x-dimension representing percent maximum reward and the y-dimension representing the percent maximum pain cost. Observed choices are overlaid points, the filled color represents the observed decision (red = reject, blue = accept). The point outline represents model-based confidence (yellow = ambivalent, purple = confident). Decision boundaries are overlaid in white. There were substantial individual differences in choice behavior and model-estimated value, as well as differences in choice consistency (indicated by the width of the band of neutral color surrounding the decision boundary). Note that decision boundaries diverged considerably from objective perceptual equality (line of reward = cost, not displayed).

### GLM 1: Parametric modulation by SV and confidence

In GLM 1 we measured parametric modulation of BOLD responses by continuous regressors for SV and choice confidence over the course of value-based choices. During the offer phase SV correlated significantly with activation in many regions of cortex, including a network of value-related regions incorporating vmPFC, posterior cingulate cortex, orbitofrontal cortex (OFC) the basal ganglia, posterior insula, and hippocampus, as well as other regions known to be involved with perceptual and value comparison such angular gyrus, lateral temporal cortex, and visual cortex (Fig 3A). A comprehensive list of significant clusters for all conditions are tabulated in S1 Table.

**Fig 3:**
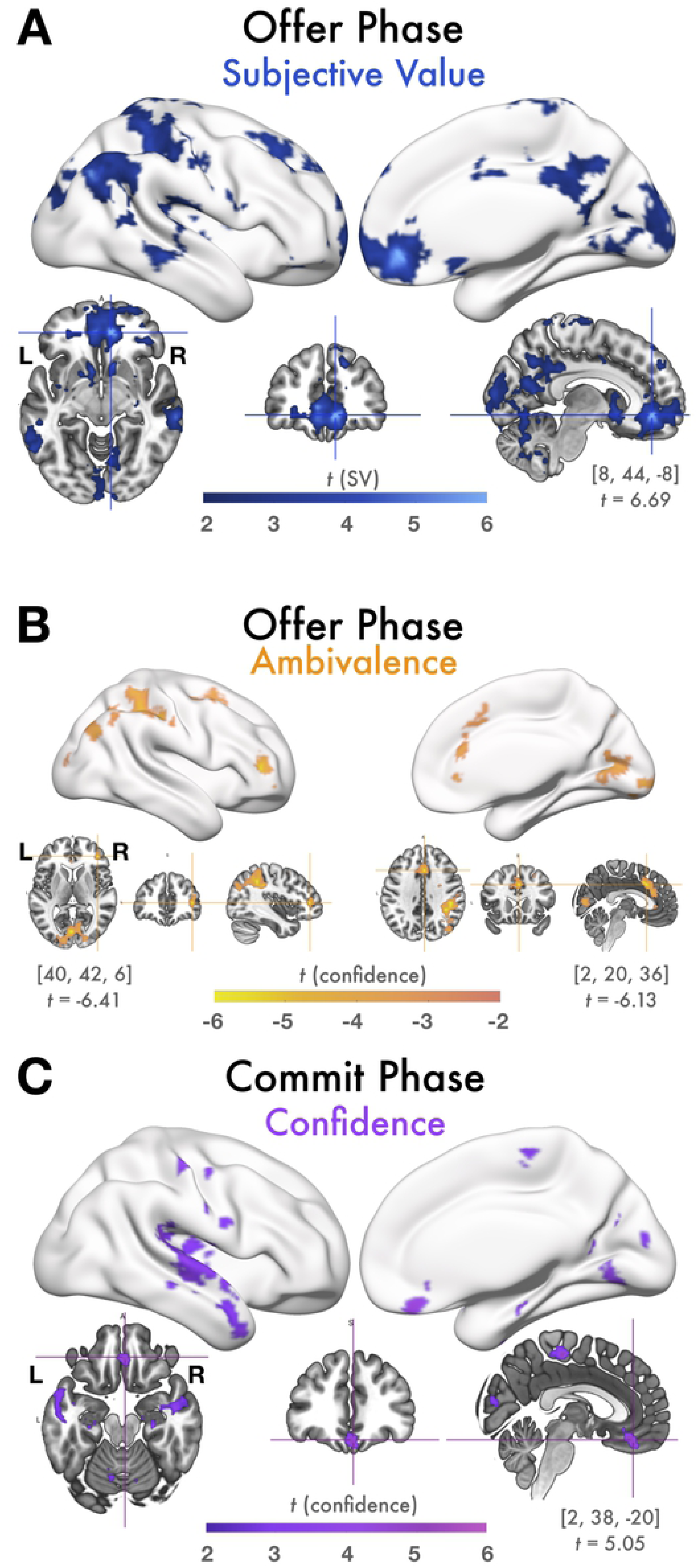
GLM1: Parametric correlation of BOLD with SV and Confidence. During the offer phase, vmPFC tracked SV but not confidence and during the commit phase, vmPFC tracks confidence but not SV. Figures display all statistically significant results at p<.05 TFCE-corrected at the whole-brain level. **(A)** BOLD responses during the offer phase that correlated positively with model-estimated SV. During the offer phase, a large cluster of voxels in vmPFC tracked SV while participants evaluated the offer. Slice images show local peak activation in vmPFC, along with the MNI coordinates and t-value of the local maximum in this cluster. There was also significant activation throughout the value network including within posterior cingulate cortex, the basal ganglia, insula, and hippocampus; regions involved with value-comparison such as angular gyrus and lateral temporal cortex; and visual cortex. No voxels correlated positively with SV during the commit phase, suggesting that value-responses, particularly in vmPFC, emerge relatively early in the decision making process. **(B)** BOLD responses positively correlated with ambivalence (inversely correlated with confidence) during the offer phase. Areas involved with cognitive control and response competition, such as lPFC and dACC tracked ambivalence while participants deliberated accepting or rejecting the offer. Slice images show local maxima in regions of theoretical interest (lPFC and dACC). No voxels in vmPFC or elsewhere responded positively with choice confidence during the offer phase, suggesting that neural responses corresponding to high confidence emerge relatively later than SV and ambivalence. **(C)** BOLD responses correlated positively with model-estimated confidence during the commit phase. During the commit phase, while participants submitted a response and viewed feedback about their decision, a cluster of voxels in vmPFC tracked decision confidence, as well as regions including posterior insula and lateral temporal cortex. Slices images show peak activation in vmPFC.

We additionally found a relatively smaller set of regions where activity negatively correlated with confidence during the offer phase including areas associated with cognitive control and conflict resolution such as dACC and right lPFC, right superior parietal lobule (SPL), premotor regions, and visual cortex (Fig 3B). There were no voxels with activity negatively related with SV nor any voxels that were positively related with confidence during the offer phase. Notably, neural responses to neutral SV can’t be easily interpreted with the SV regressor alone. Instead, the inverse of the confidence regressor can be interpreted as a measure of choice conflict. Both the dACC and lPFC demonstrated a strong inverse relationship with confidence, consistent with previous results demonstrating their recruitment during ambivalence, conflict, uncertainty, and choice difficulty during value based decision making and other tasks (21,46–52) . The inverse relationship between the confidence regressor and conflict-resolution and positive relationship of the regressor with SV is compatible supports the idea that neural activity during the offer phase is primarily related to valuation processes.

Surprisingly, we did not observe any BOLD responses, in vmPFC or elsewhere, that varied with SV during the commit phase. Furthermore, during the commit phase, there were no regions where activity varied negatively with choice confidence. Instead, choice confidence significantly predicted activity in a cluster of voxels within vmPFC as well as other regions including posterior insula, superior temporal cortex, and premotor areas during the commit phase (Fig 3C).

To characterize the dynamic relationship between SV and confidence signals in the vmPFC, we performed a FIR model. As shown in Fig 4 beginning ∼ 3 seconds after the presentation of an offer (consistent with a response delayed by the HRF), there is a rapid increase of SV related activity in vmPFC lasting approximately seven seconds. In contrast, activity related to confidence emerges approximately 10 seconds after the offer (which is 5.5 seconds after the presentation of the response cue). As this activity reaches a peak around 13 seconds, SV related activity has already returned to baseline, demonstrating a clear cross-over in the timing of these two variables.

**Fig 4:**
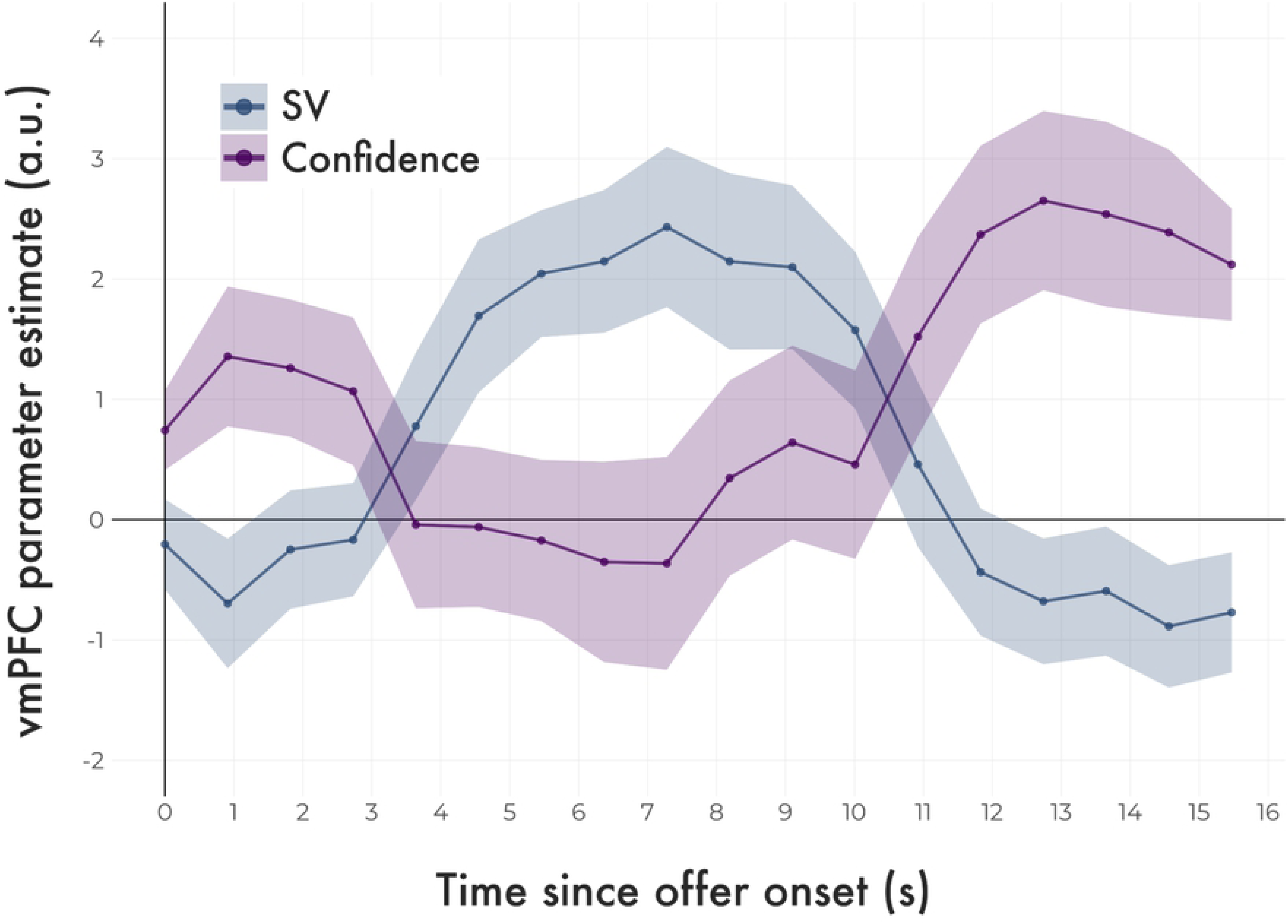
Time course of vmPFC responses tracking SV and Confidence. SV signals preceded confidence signals in vmPFC. Neural responses were sampled once per TR (every 910 ms), beginning at the onset of the trial stimulus, marked here as 0 s. Plotted lines (group means) and ribbons (SEM) represent vmPFC parameter estimates over the course of the trial, estimated with an FIR model. The blue line traces the strength of the association between BOLD responses in vmPFC and model-estimated SV, the purple line traces their association with model-estimated confidence.

Taken together, the key finding from GLM1, whether estimated by classic HRF convolution or FIR methodologies was that during the offer phase, vmPFC tracks SV but not confidence, and during the commit phase vmPFC tracks confidence but not SV. These results suggest that in value-based decision making, vmPFC responses do increase with both SV and confidence, however these signals do not modulate simultaneously. Rather, there may be a more dynamic process through which vmPFC is first involved in the valuation process before transitioning to signaling confidence about that valuation. This interpretation supports the idea that vmPFC’s dual roles in value-based decision making are separable if that process is examined in sequential stages.

### GLM 2: Categorical choice by confidence conditions

While SV predicts choice outcomes (i.e. offers perceived as highly valuable are likely to be accepted), SV and outcome are not perfectly related, especially when choices are closer to one’s decision boundary. It is possible that SV is computed relatively early on, but that vmPFC responses relating to the final choice (selected with respect to SV) occur later and concurrently with choice confidence, during the commit phase. Therefore, in GLM2 we measured responses related to choice outcomes and confidence. To do this, GLM2 modeled BOLD responses from four trial conditions: AccCon, RejCon, AccAmb, and RejAmb. These analyses mirror GLM1 except that in GLM2, value is signified as trial by trial choice outcome rather than estimated SV and confidence has been binned into high (confident) and low (ambivalent) categories. During the offer phase, we observed that a similar network of regions to those that were parametrically modulated by SV in GLM1 also showed significant contrasts for decision outcome in GLM2 such that BOLD responses were stronger preceding decisions to accept than decisions to reject (Fig 5A). This augments our findings from GLM1, demonstrating that not only SV but also a decision variable is represented within vmPFC relatively early in the decision making process, in this case during the offer phase.

**Fig 5:**
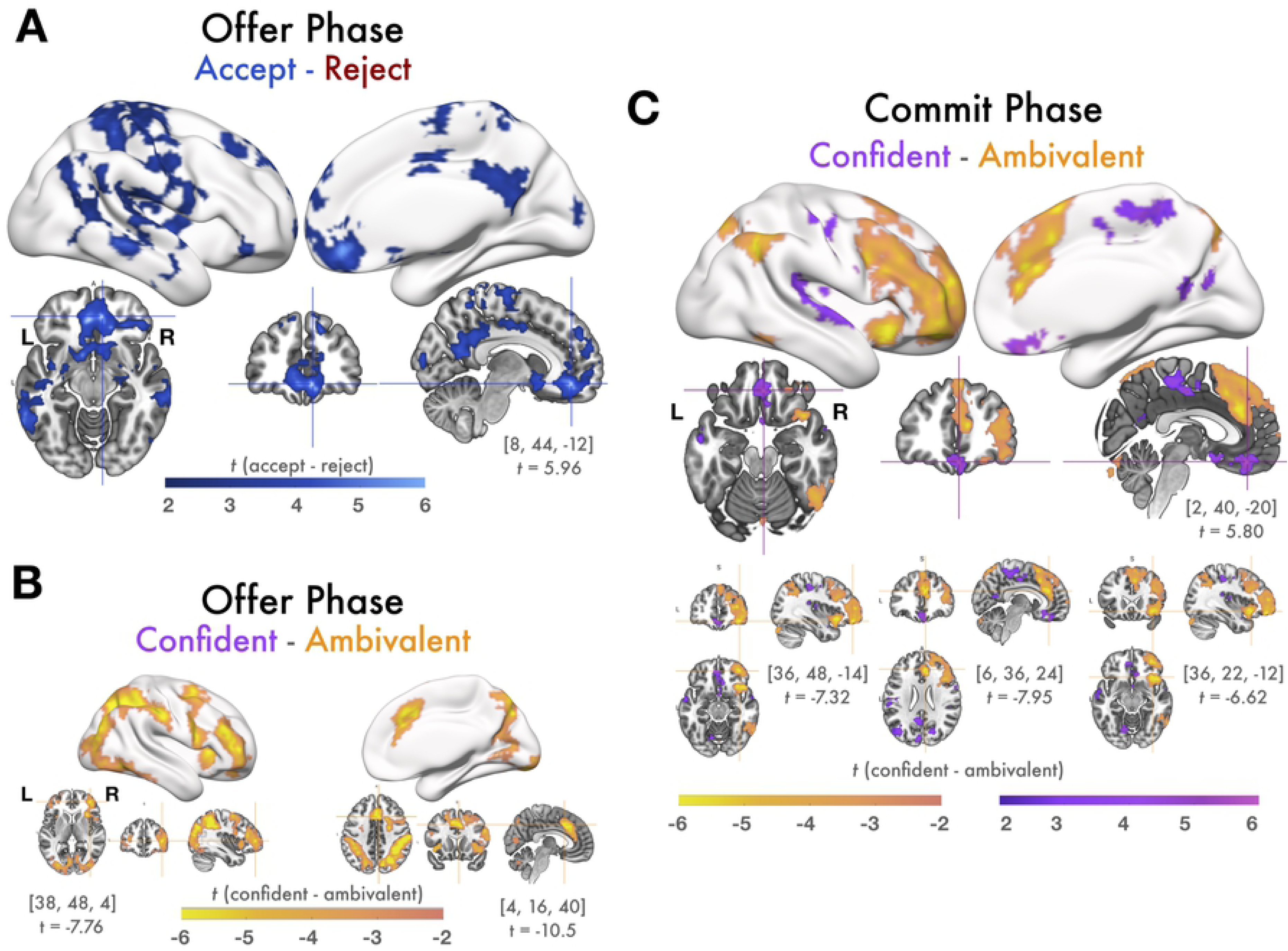
GLM2: Contrasts of decisions and confidence conditions. vmPFC encodes decision variables during the offer phase but not the commit phase, and encodes decision confidence during the commit phase, but not the offer phase. Figures display all statistically significant results at p<.05 TFCE-corrected at the whole-brain level. **(A)** Contrasts of response magnitudes from the offer phase of accept trials > reject trials: During the offer phase, clusters of voxels in vmPFC and other regions in the value network, many of which were also parametrically correlated with SV, had stronger response magnitudes preceding accept decisions than reject decisions. Slices images show peak coordinates in vmPFC. No voxels exhibited this contrast during the commit phase, nor were there any regions with stronger responses during rejected trials than accepted trials during either phase. **(B)** Contrasts of response magnitudes from the offer phase of confident trials < ambivalent trials. Slice images illustrate peak activation in lPFC and dACC. **(C)** Contrasts of response magnitudes from the commit phase of confident > ambivalent trials (purple) were observed in vmPFC were stronger for confident trials than ambivalent trials. No regions showed the same contrast during the offer phase. Clusters of voxels in lateral OFC, the anterior insula, and dACC showed the reverse pattern, with stronger response magnitudes during the commit phase of ambivalent trials than confident trials. Note that, several regions (dACC, lPFC, lateral OFC, and anterior insula) show the confident < ambivalent contrast during both the offer and commit phases. Slice images show peak coordinates for confident > ambivalent in vmPFC and peak coordinates for confident < ambivalent in lateral OFC, dACC, and anterior insula.

There was substantial overlap between value-related responses in GLM1 and decisions to accept in GLM2. These similarities are likely attributable to participants’ relatively stable choice behavior, causing SV model estimates to be strongly predictive of choice outcomes. Consequently, neural responses associated with high estimated SV in GLM1 were also categorized as objective accept choices in GLM2. The primary exception to this rule was visual cortex, where activity tracked SV but not choice outcome, suggesting that visual cortex activation varied with SV only insofar as it systematically related to visual properties of the offer stimuli.

Similarly, the contrast of BOLD responses from the offer phase of ambivalent versus confident choices revealed a similar network of regions as those that were negatively related to confidence during the offer phase from GLM1, with the addition of the anterior insula for GLM2 (Fig 5B). No regions, including vmPFC, showed a stronger response during confident choices than ambivalent choices during the offer phase, nor were any regions more responsive during rejected trials than accepted trials.

During the commit phase, there were highly significant and widespread differences between confident and ambivalent choices, but no regions responded preferentially to one choice outcome (accept or reject) over the other. Critically, clusters of voxels in vmPFC as well as posterior cingulate, and an adjacent medial segment of superior parietal lobule showed a significant contrast between confident and ambivalent decisions during the commit phase, with stronger BOLD responses to confident choices (Fig 5C, purple). The reverse contrast (ambivalent versus confident) revealed several of the same regions that responded selectively to ambivalent choices during the offer phase, such as the anterior insula, lateral PFC, and dACC (Fig 5C, orange). This could possibly indicate sustained ambivalence-related activity that arises early in the valuation process and may still be unresolved at the time of choice commitment for highly difficult choices. The observed responses in vmPFC, selective for positive choice outcomes during the offer phase and for high confidence during the commit phase, provide additional evidence of separable stages of activity over the course of a value-based choice.

Given the robust SV-(from GLM1) and choice-related (from GLM2) responses during the offer phase, it is possible that apparent confidence-related signals in vmPFC activity during the commit phase were in fact attributable to strong, sustained value-related responses carrying over from the offer phase. That is, relative differences in vmPFC responses to AccCon choices versus AccAmb choices could have driven the observed main effect of confidence – and moreover these effects might be better explained by value than confidence. To verify that the apparent effects of confidence could not be explained by relative differences in value, we measured contrasts of confident versus ambivalent BOLD responses separately for accepted and rejected offers. We were specifically interested in these comparisons within our a priori vmPFC ROI and therefore restricted statistical correction to this region alone (Fig 6A). During the commit phase, we observed a cluster of voxels with significantly stronger responses during RejCon versus RejAmb trials as well as a cluster of voxels that preferred AccCon to AccAmb (Fig 6B). Notably, because SV of RejCon trials is less than SV of RejAmb trials, this result suggests that during the commit phase, clusters of vmPFC activity are signaling choice confidence irrespective of SV. We found no voxels that demonstrated the same pattern during the offer phase, nor did we find any voxels with stronger responses to ambivalent choices during either phase.

**Fig 6:**
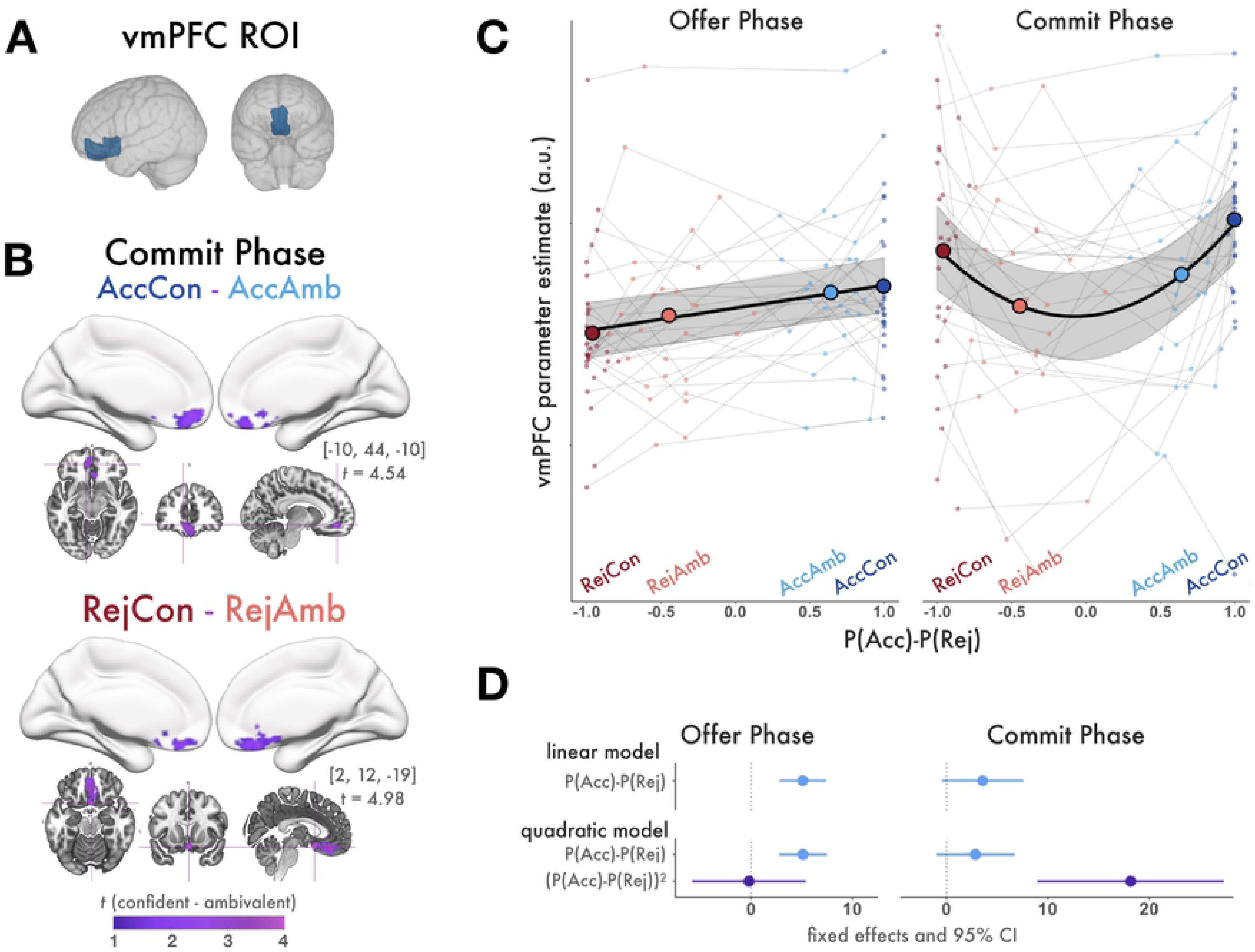
vmPFC ROI analysis. **(A)** Anatomical vmPFC ROI **(B)** AccCon > AccAmb and RejCon > RejAmb contrasts of BOLD responses during the commit phase. Figures display all statistically significant results at p<.05 TFCE-corrected within the vmPFC ROI. During the commit phase, vmPFC responses are stronger for high confidence choices, even for reject choices when confidence is inversely related with SV. **(C)** vmPFC Parameter estimates from the offer (left) and commit phase (right). Small points are individuals’ raw vmPFC parameter estimates with respect to their mean P(Acc)-P(Rej) of each trial condition. Large points are group mean vmPFC parameter estimates with respect to the group mean P(Acc)-P(Rej) of each condition. The fit line indicates regression predictions and 95% CI from the winning model for that decision phase. During the offer phase, vmPFC responses increased linearly across trial conditions with increasing value. During the commit phase, they increased quadratically. **(D)** Comparison of mixed effects regression models predicting vmPFC parameter estimates (GLM2) from the and quadratic (bottom) extensions of value. Model estimates and 95% confidence intervals show fixed effects of linear and quadratic model terms, confidence intervals not spanning 0 are considered significant. vmPFC parameter estimates from the offer phase (left) were best fit with a linear model, indicating that during the offer phase, vmPFC tracks value. vmPFC parameter estimates from the commit phase (right) were best fit with a quadratic model, indicating that during the commit phase vmPFC tracks confidence.

### Decision phase-specific responses in vmPFC

Finally, for both phases of the decision making process, we examined the pattern of response magnitudes in vmPFC across trial conditions of increasing value, beyond relative differences between pairs of conditions and compared the fits of models with linear and quadratic terms (Fig 6C). For the offer phase, the linear model best explained the observed pattern of vmPFC parameter estimates across the four conditions as the additional term in the quadratic model did not improve the model fit (Linear model: AIC = 901.1, BIC = 911.98; Quadratic model: AIC = 903.1, BIC = 916.69; comparison of models: χ2(1) = .0018, p = .966), indicating that vmPFC activity increased linearly with P(Acc)-P(Rej) during the offer phase (signaling value). For the commit phase, the quadratic model best explained the observed pattern of vmPFC parameter estimates (Linear model: AIC = 1014.5, BIC = 1025.3, Quadratic Model: AIC = 1002, BIC = 1015.6, comparison of models: χ2(1) = 14.320, p < .001), indicating that vmPFC responses take a quadratic function with respect to value during the commit phase, signaling confidence. Model effect sizes are plotted in Fig 6D and details about model fits are shown in Table 1.

In summary, vmPFC responses increased linearly across trial conditions of increasing value during the offer phase, signifying early involvement with valuation of the offer stimulus, and varied quadratically across the same conditions during the commit phase, signifying late involvement with valuation of the decision and deriving confidence. Notably, no values associated with model-estimated confidence (such as those used in GLM1) were entered into the mixed effects models. Rather, the models estimated participants’ mean parameter estimates from the four trial conditions (of GLM2) from their mean P(Acc)-P(Rej) for all trials of that condition. Therefore, the plots in Fig 6c demonstrate that during the offer phase, vmPFC responses naturally take a linear function across the four trial conditions of ascending value P(Acc)-P(Rej), whereas during the commit phase, vmPFC responses naturally take a quadratic function across the same trial conditions.

Strictly speaking, because P(Acc)-P(Rej) takes a sigmoid function with respect to SV and that vmPFC increased linearly with SV in GLM1, one might predict vmPFC responses to take a logistic function with respect to P(Acc)-P(Rej). Visual inspection of individual raw vmPFC parameter estimates (Fig 6B) did not reveal that the underlying pattern of vmPFC parameter estimates had a logistic shape. To verify that this was the case, we repeated the linear mixed regression model, substituting SV predictors for P(Acc)-P(Rej) predictors, and found analogous statistical results. While the fMRI parameter estimates had sufficient resolution to discern between linear and quadratic functions, it is unlikely that a similar distinction could be made between two monotonic functions such as linear and logistic functions. We present results with respect to P(Acc)-P(Rej) because this measure improved the interpretability of group-wide results by controlling the range of predictor values across the group while preserving the sign of raw SV (as described above).

#### Behavioral RTs and Confidence

Before analysis, trials with RTs exceeding 3 SD of the overall group mean were excluded (1.56%). Additionally, trials exceeding 3 SD of the individual’s mean RT for each bin of each analysis were excluded (0.98%). The model confirmed that RTs times took an inverse quadratic function with respect to value (Fixed Effects: (P(Acc)-P(Rej) estimate = -35.415, 95CI [-50.042, -20.787], SE = 7.466, t = -4.743, p < .001; P(Acc)-P(Rej)^2^ estimate = -77.330, 95CI [-102.213, -52.451], SE = 12.700, t = -6.90, p <. 001; Random Effect SD = 88.02), indicating that participants were slower to commit to decisions associated with low model-estimated confidence (Fig 7). Notably, in addition to the quadratic relationship, there was also a negative linear correlation between value and RTs, indicating that it took participants longer to commit to decisions about low value offers versus high value offers. This may have been caused by the observed bias to accept offers more offers than were rejected, which meant that responses to lower value offers, which were often reject choices, required overriding the default decision to accept.

**Fig 7:**
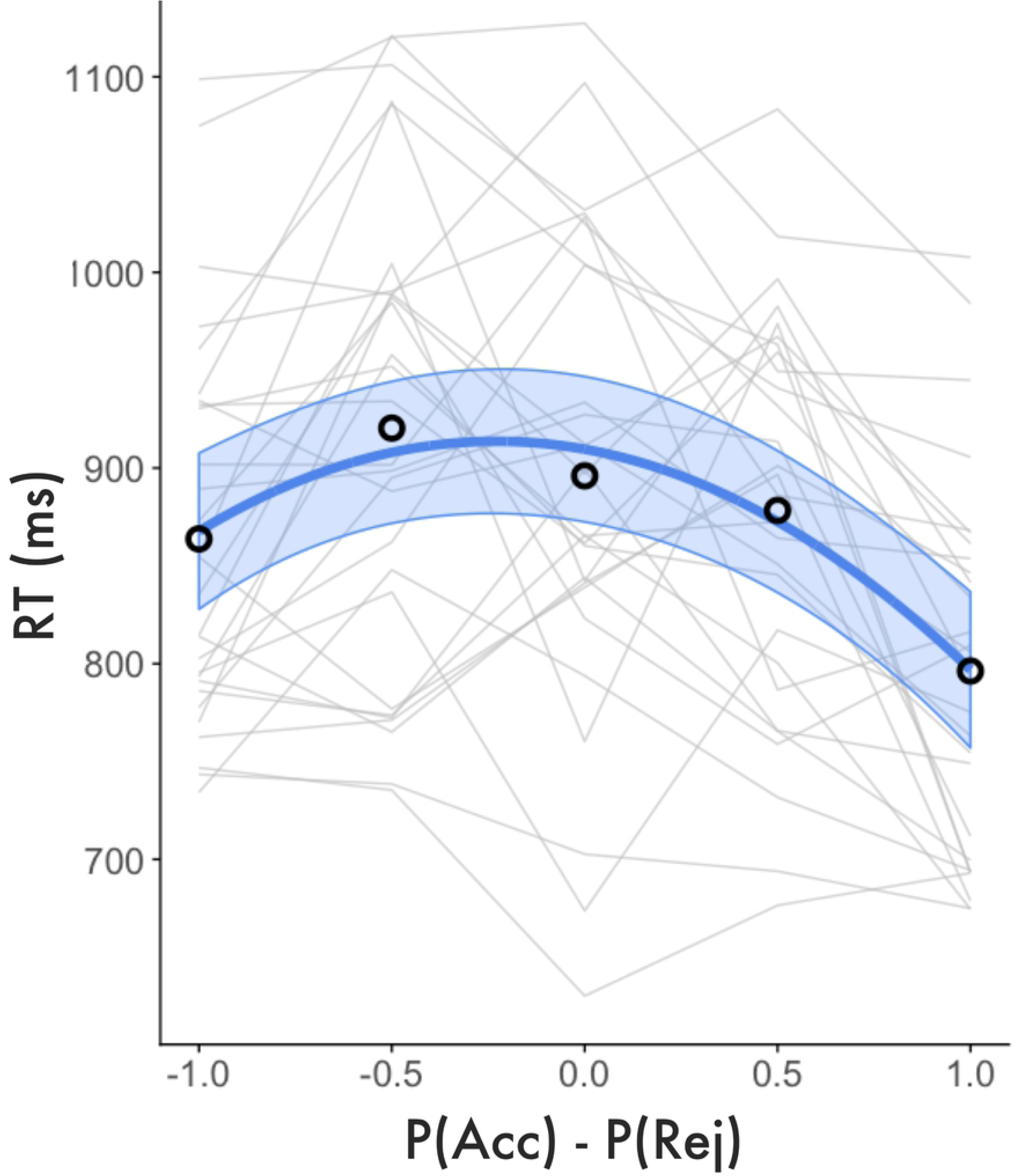
RT analysis. Individual and group mean RTs (small lines) are plotted with respect to value (P(Acc)-P(Rej)). Fit lines represent model predicted RTs and 95% CIs. RTs take an inverse quadratic function with respect to model-estimated value. The time it took participants to commit to decisions was negatively correlated with model-estimated confidence (the quadratic extension of value).

We observed statistically analogous, but somewhat weaker results when using SV as value predictors (Fixed Effects: SV estimate = -27.198, 95CI [-32.326, -22.070], SE = 2.618, t = -10.390, p < .001; SV^2^ estimate = -7.900, 95CI [-12.230, -3.563], SE = 2.212, t = -3.570, p = .001; Random Effect SD=8.020), which is likely due to large individual differences in estimated SV, as described above. Taken together, the results of the RT analysis confirmed that there was a meaningful relationship between model-estimated confidence and behavior. Participants were slower to commit to decisions about offers we predicted would elicit ambivalence, suggesting that these choices might have been more challenging to resolve. This is not an indication that RTs measure the subjective experience of confidence, but rather that there is an empirical basis that model-estimated confidence captures aspects of our task associated with decision difficulty. Importantly, because participants varied in their subjective preferences and their decision boundaries diverged considerably from the reward=cost line (Fig 2), increased RTs for ambivalent choices cannot be explained by difficulty in perceptual discrimination (i.e. merely determining whether costs or rewards were perceptually larger).

## Discussion

We measured behavioral and neural responses during a two-phase Ap-Av decision making task with consequential mixed outcomes. Model-based SV, confidence estimates and decision variables correlated with BOLD responses throughout the cortex, including within vmPFC. Our series of results provides strong support for the recent suggestion that vmPFC encodes both value and confidence during value-based decision making (17,19,21). Importantly, the present research is the first evidence of this phenomena in the realistic context of deterministic approach avoidance choice scenarios. It extends previous findings from studies that have used different approaches for asessing confidence, including explicit rating of hypothetical SV for items or events (17), deciding between two positively-items (19), or decisions with probabilistic risk (21). We then presented novel evidence that such vmPFC signals emerge during different phases of decision making. Specifically, neural responses to offer valuation (SV) and choice determination were temporally dissociated from decision valuation (confidence), with the former dominating vmPFC prior to choice commitment, when the latter takes priority. The following sections discuss our key findings regarding neural signatures of SV and confidence as well limitations of this study and suggestions for future research.

Our first finding was new evidence of approach-selective vmPFC responses preceding choice commitment in an economic ApAv task. During the offer phase of each trial, participants deliberated on accepting or rejecting offers and vmPFC responses increased with estimated SV (GLM1). We complemented this finding by demonstrating that parameter estimates from an anatomical vmPFC ROI increased monotonically with P(Acc)-P(Rej) (GLM2), consistent with a vast literature documenting vmPFC’s role in integrating items’ cost and reward attributes into its overall SV (4,11,12,53). vmPFC also distinguished between decision outcomes during the offer phase, with stronger responses anticipating accept decisions than reject decisions (GLM2). These results may be partially explained by goods-based models of value-based decision making, which propose that economic choices are made between goods rather than actions (54), and can therefore be settled prior to planning the action to submit the decision (55).

Remarkably, we found no regional activation that was significantly modulated by SV (GLM1) or that differed by choice outcome (GLM2) during the commit phase, even when statistical correction was restricted to vmPFC (GLM2). This result was unexpected given previous findings that vmPFC representations of subjective preferences are automatically elicited by task stimuli, including contexts in which such information is task-irrelevant (7, 56). Moreover, SV responses in vmPFC can sustain the entire presentation of value stimuli with durations far longer than the time needed to make a decision, regardless of whether the additional time affects decision behavior (57). Together, these prior findings suggested that in our study, vmPFC would encode SV whenever the offer stimuli were visible, including during the commit phase. Our failure to find such effects may indicate that a two-phase decision task, such as the one we implemented, reveals the transition of vmPFC processing from offer valuation to decision valuation in such a way that other paradigms that either require immediate responses, only briefly present value stimuli, or that rapidly transition decision outcomes after choice commitment do not.

One alternative hypothesis is that there were in fact enduring but relatively weak SV representations in vmPFC during the commit phase that did not reach statistical significance. Visual inspection of the group mean vmPFC parameter estimates during the commit phase (GLM2) suggests a somewhat asymmetrical quadratic function such that accepted offers, on average, were associated with larger response magnitudes than rejected offers. However, a linear model provided only a marginal fit to the same data, consistent with only a subtle positive correlation, if any, between vmPFC activation and value during the commit phase. Thus, there is a hint that vmPFC represents SV, albeit weakly, through the commit phase, but this effect is markedly exceeded by confidence.

There have been various empirical and theoretical suggestions that confidence signals emerge early in decision making, evolving in parallel with processing of choice stimuli and the decision itself (17,22,23,58). Others have reported evidence that confidence lags behind decision variables and value estimates, emerging closer to the time of choice commitment or even later (25,27–32). We presented a series of results that converged to decidedly endorse the latter, and to our knowledge provide the first evidence for delayed confidence processing in the context of value-based decision making. Specifically, we did not observe any BOLD responses that were parametrically correlated with confidence during the offer phase (GLM1) nor were there any regions with responses during the offer phase that were selective for confident over ambivalent trials (GLM2), even when statistical correction was restricted to vmPFC. Confidence responses were observed after responses related to the decision information itself, the SV of the offer and the decision outcome. During the commit phase of each trial, while participants submitted a choice and then viewed feedback about their selection, vmPFC responses increased parametrically with decision confidence (GLM1) and similarly were stronger for confident choices than ambivalent choices (GLM2). Closer inspection revealed that vmPFC parameter estimates from the commit phase took a quadratic function with respect to increasing value (GLM2). Therefore, during the commit phase, vmPFC responses were stronger for confident choices than ambivalent choices, even for rejected offers when high confidence is associated with low SV. Visualization of the time course SV and confidence responses in vmPFC (GLM1 FIR) illustrated a distinct transition from early value-related signal to late confidence-related signal in vmPFC. It is noteworthy that the SV and confidence regressors peaked at similar delays following the onset of the offer stimulus and response cue, respectively. Although the delay of the BOLD response makes it difficult to identify precisely when in the trial this transition occurred, this may be preliminary evidence that the most robust confidence-related response was triggered by the cue for the participant to commit a response.

Beyond vmPFC, we observed extensive overlap between regions tracking SV during the offer phase (GLM1) and regions that selectively activated preceding accept decisions (GLM2). This was observed in a larger value-network including regions important for sensory association, value-comparison, and reward processing such as angular gyrus, posterior parietal cortex, lateral temporal cortex, posterior cingulate cortex, and the ventral striatum. The value network likely encodes finer-grained distinctions between SV and choice outcome than were apparent in our analyses. For example, others have suggested separable time courses for evolving value signals and decision variables (59). Our paradigm was well suited for direct comparison of SV and confidence signals but not necessarily for dissociating SV from decision variables, given the strong correlation between the two. Future research aiming to find finer dissociations of SV and decision variables may benefit from novel variations of our task.

Furthermore, our anatomical vmPFC ROI was selected a priori and is somewhat inclusive, combining subcallosal cortex and medial frontal cortex from the Harvard-Oxford atlas, which include structures that others have labeled medial OFC, pregenual or subgenual ACC. Thus, we cannot draw meaningful conclusions about functional specificity at a smaller scale but we appreciate that others have made interesting discoveries on this front. For example, it has been suggested that over the course of stimulus processing, OFC is the first cortical site where appetitive stimuli are assigned reward, whereas later processing in vmPFC transforms value representations into choices (53) that guide action selection (60). A previous fMRI study that employed a strategy game requiring choices to attack or defend reported that an adjacent region, rostral ACC, preferentially responded when participants defended versus attacked and tracked the value of deploying defense strategies but not attack strategies (10). While we did not observe regional activation that correlated negatively with SV nor regions that selectively responded to reject choices, the conceptual equivalent to defending, the notion of separable but adjacent neural bases for approach and avoidance behaviors is compelling. Combining finer parcellation of prefrontal cortex with fMRI models that include terms for individual offer attributes reveal separable patterns of responses to reward, SV, and choice outcome may be an interesting avenue for future research. However, this was beyond the scope of our study.

Others have demonstrated that model-based confidence estimates correspond closely to self-reported confidence (17) and the strength of the relationship between model-based confidence and vmPFC activation strongly predicts the relationship between self-reported confidence and vmPFC activation (15). A key aspect of our study was the characterization of confidence without a reliance on post-hoc self-report or metacognition. Using a naturalistic task and implicit measures of confidence, we replicated previous findings that model-estimated confidence had an inverse quadratic relationship with RTs (17), such that increasing model-estimated confidence predicted faster decision commitments (Fig 7), suggesting that model-estimated confidence captures an element of choice difficulty.

A similar study by De Martino et al. (2013), that incorporated explicit confidence ratings with an fMRI value-based choice task suggests separable neural signals corresponding to model-based and self-reported confidence and found that while vmPFC tracked unsigned value differences between choice options (which roughly corresponds to our model-estimated confidence), rostrolateral PFC (rlPFC) tracked self-reported confidence. They used this basis to ground a hypothesis that rlPFC probes internal confidence signals, represented in vmPFC, and makes them available for metacognitive self-report. Others have made similar claims (61). Notably, rlPFC responses correlated with ambivalence in our task (which did not require explicit confidence ratings), suggesting that our task did not elicit similar metacognitive appraisal processes. While there is not yet broad consensus regarding the loci of implicit versus metacognitive confidence – it seems evident that decision tasks requiring metacognitive confidence judgments may recruit unique neural resources from tasks that model confidence directly from decision behavior. Nonetheless, our results support recent suggestions that implicit confidence, the valuation of one’s judgment, shares a neuroanatomical basis with a variety of other valuation processes, vmPFC (17,20,30). We add to this growing theoretical framework evidence for a dynamic process by which vmPFC shifts from valuation of external stimuli to valuation of internal value representations and the decisions they inspire.

## Acknowledgements

This work supported by Grant W911NF-16-1-0474, Contract W911NF-09-0001 and Cooperative Agreement W911NF-19-2-0026 with the Army Research Office of the Army Research Laboratory. We thank Mario Mendoza, Gold Okafor, and Viktoriya Babenko for their assistance with data collection as well as Neil Dundon, Michelle Marneweck, Tyler Santander, and Evan Layher for helpful suggestions.

## Supporting Information Captions

**S1 table: Significant cluster activation from GLM1 and GLM2**

Full details of significant task-related activation. Clusters <10 voxels excluded. The first three columns identify the analysis, decision phase when BOLD responses were measured, and the variable measured in the statistic (Con. = confiden(t)ce, Amb.=ambivalen(t)ce. The next column specifies each cluster and indicates the total number of voxels in that cluster. The remaining columns are details about local maxima within that cluster, including t-value (* signifies cluster peaks illustrated in figures), MNI coordinates, region from AAL of MNI space, and Brodmann area (-signifies peak voxel falls outside of labelled cortex, region provided is the closest).

